# A Bistable Mechanism Mediated by Integrins Controls Mechanotaxis of Leukocytes

**DOI:** 10.1101/509091

**Authors:** Alexander Hornung, Thomas Sbarrato, Nicolas Garcia-Seyda, Laurene Aoun, Xuan Luo, Martine Biarnes-Pelicot, Olivier Theodoly, Marie-Pierre Valignat

## Abstract

The recruitment of leukocytes from blood vessels to inflamed zones is guided by biochemical and mechanical stimuli, with mechanisms only partially deciphered. We studied here the guidance by flow of primary human effector T lymphocytes crawling on substrates coated with ligands of integrins LFA-1 (α_L_β_2_) and VLA-4 (α_4_β_1_), and showed that cells segregated in two populations of opposite orientation for combined adhesion. Sharp decisions of orientation were shown to rely on a bistable mechanism between LFA-1-mediated upstream and VLA-4-dominant downstream phenotypes. At the molecular level, bistability results from a differential front-rear polarization of both integrins affinity, combined with an inhibiting crosstalk of LFA-1 toward VLA-4. At the cellular level, directivity with or against the flow is mechanically mediated by the passive orientation of detached uropod or lamellipod by flow. This complete chain of logical events provides a unique mechanistic picture of a guiding mechanism, from stimuli to cell orientation.

**Significance:** Cellular guidance is crucial to many biological functions, but the precise mechanisms remain unclear. We have analyzed here an original phenotype of flow-guided cells mimicking leukocytes crawling into the blood vessels and showed that the *controlling parameter* of cells decision to migrate upstream or downstream was the *relative number* of two specific adhesion molecules, the integrins LFA-1 and VLA-4. The spatial polarisation of integrins affinity and an intermutually feedback of their activation create a bistable system where cells adhere either by their tip or their tail and orient respectively downstream or upstream. This mechanism therefore proposes a complete chain of event from stimuli to cell orientation and differs strongly from the chemotaxis paradigm because stimuli trigger no signaling.

## INTRODUCTION

Cell guiding is involved in numerous essential functions of living organisms, however its mechanisms remain partially understood. Chemical guiding, or chemotaxis, has long been studied and many mechanistic elements have been identified^1,2^. In contrast, mechanical guiding, or mechanotaxis, has been acknowledged more recently, and although many physiological roles have been identified in differentiation^3^, morphogenesis^4,5^, or leukocyte activation^6^, its basic functioning remains largely open. The regulation of immune cells trafficking between lymphoid organs, blood system, and inflamed or infected zones involves robust guiding mechanisms by chemical signals^7–9^ and also mechanical signals like hydrodynamic shear stress^10–20^. The detection of external force by leukocytes arguably relies on integrins adhesion proteins^21^, which undergo conformational changes by inside-out ^22,23^ and outside-in^24,25^ signaling, and transduce intracellular signaling when submitted to force. Integrins, which are already key players in the recruitment of leukocytes from blood flow ^18,26–28^, may also orchestrate leukocytes mechanotaxis under flow. A precise mechanism of mechanotaxis controlled by integrins is however lacking.

Both *in vivo* and *in vitro* experiments have reported guiding by flow of leukocytes crawling on the internal walls of blood vessels. *In vivo*, on a rat model suffering from autoimmune encephalomyelitis, effector T cells crawl preferentially upstream on the luminal surface of leptomeningeal vessels presenting ICAM-1 and VCAM-1^10^, ligands of LFA-1 and VLA-4 integrins respectively. Mouse T cells on monolayers of blood brain barrier endothelial cell cultured *in vitro* were also crawling upstream in an ICAM-1 required manner ^13^. On acellular glass substrates, human effector T lymphocytes on ICAM-1^12–14^ and mouse T-cells on ICAM-1 and ICAM-2 crawled upstream^19^, whereas neutrophils and metastatic lymphocytes crawled downstream on ICAM-1^11,12,19^ and on VCAM-1 ^14,29^. This variety of mechanotaxis responses versus the types of leucocytes and of integrins suggests the existence of sophisticated mechanosensing mechanisms controlled by integrins.

Two types of mechanisms of orientation by flow have been proposed for leukocyte adhering with integrins. A first approach proposes an “active” mechanism inspired from the chemotaxis machinery. Cue detection (shear stress direction) involves outside-in signaling at anchoring sites mediated by integrins functioning as molecular force transducers ^30–33^. Evidences of integrin-based signaling during migration under flow have been reported for endothelial cells ^33,34^ and neutrophils ^21^. Such mechanisms are “active” in the sense that cells develop a specific intracellular signaling activity in response to flow and rely therefore on mechanotransduction. Alternatively, a “passive” model was established for upstream crawling T lymphocytes^13^. In this model, flow direction is detected by the passive orientation of cell tail (or uropod), which is not adherent and rotates in a flow like a wind vane in a breeze. Reorientation of the whole cell against flow follows tail orientation via the re-alignment of cell’s front by the on-going mechanism of front-rear polarization. This mechanism is “passive” in the sense that it requires no signaling triggered by the external cue and therefore no mechanotransduction. Whether the active and passive mechanisms are specific to certain cells types, or whether they are alternatively triggered by different microenvironments remains unclear.

A fundamental question in leukocyte mechanotaxis concerns therefore the functional role of integrins in terms of adhesion mediators and/or mechanotransducers. The fact that the mechanotaxis phenotype of T cells changes from upstream to downstream when the substrates are coated by ligands of integrins LFA-1 or VLA-4 ^14,19,29^ raises fundamental questions such as what decides upstream versus downstream guiding of leukocytes, or how different integrins control different orientation. In this paper, we analyzed T lymphocyte migration on substrates with quantified mixtures of ICAM-1 and VCAM-1 molecules. We show that a bistable mechanism triggers upstream and downstream mechanotaxis phenotypes of T cells on mixed ICAM-1/VCAM-1 substrates, and that bistabilty relies on a combination of molecular and cellular mechanisms. At molecular level, a crosstalk between integrins LFA-1 and VLA-4 and the contrary polarization of LFA-1 and VLA-4 affinity sustain a differential adhesion of cells either by their leading or trailing edge. At the cellular level, polarized adhesion triggers passively the upstream phenotype by a mechanism of wind vane/uropod or the downstream phenotype by a mechanism of lamellipod flow focusing. This bistable model presents a complete functioning of mechanotaxis controlled by integrins, where the logical chain of mechanistic elements are identified from molecular to cellular level and from stimulus to cell orientation outcome.

## RESULTS

### Flow fosters a variety of migration phenotypes on mixtures of ICAM-1 and VCAM-1

To examine the respective role of LFA-1 and VLA-4 on the orientation of T lymphocyte crawling under flow, we prepared surfaces coated with either ICAM-1, VCAM-1 or mixtures of ICAM-1 and VCAM-1 by adsorption of Fc-ICAM-1 and Fc-VCAM-1 molecules into channels pre-coated with protein-A. The absolute amount of each ligand on surfaces was quantified by fluorescence (Supplementary Information 1). Without flow, the percentage of migrating cells and their speed decreased only moderately when the fraction of ICAM-1 decreased (Figure 1a,b). These observations suggest that the crawling machinery does not significantly depend on the type of integrin engaged with the substrate. In contrast, when flow was applied, cells exhibited different responses depending on the surface composition. They migrated mainly upstream on ICAM-1-treated substrates and mainly downstream on VCAM-1-treated substrates (Supplementary Movie 1), as previously reported in the literature^12,14,19^. Furthermore, the average orientation of T cells on mixed ICAM-1/VCAM-1 substrates described by the migration index on Figure 1c showed a smooth transition from upstream to downstream phenotype versus an increasing fraction in VCAM-1^8^. Flow revealed therefore a critical interplay between integrin-mediated adhesion and crawling machinery. This analysis based on population-average data is however missing important features of the system, which is in fact multiphasic. Individual cells presented indeed several distinct phenotypes for a given set of flow rate and of substrate coating. On mixed ICAM-1/VCAM-1 substrates, different T cells crawled either upstream or downstream, or rolled downstream (Figure 1d, Supplementary Movie 2). To avoid population-averaging bias, we defined precise criteria to sort cells depending on their adhesion/migration phenotype under flow. Cells were called “Non adherent” when they were flushed away by a gentle flow of 0.5 dyn.cm^-2^ after 10 min of incubation on the adhesive substrates, “Immobile” if they moved less than 30 μm during the whole acquisition (100 frames, 17 min), and “Rolling” when i-their direction remained within 10 degrees of flow direction in a 17 min path and ii-the standard deviation of their direction upon 20 s steps remained within 25 degrees. In the latter case, only the cells that rolled at velocity lower than 90 μm.min^-1^ could be tracked due to frame acquisition rate (Figure 1e, black trajectories). Finally, all remaining cells were considered as “crawling” (at this point of investigation) and sorted into upstream (Figure 1e, blue trajectories) and downstream crawling (Figure 1e, red trajectories). This description shows clearly that the transition on mixed substrates from upstream to downstream crawling phenotype is not homogeneous but biphasic.

**Figure 1:**
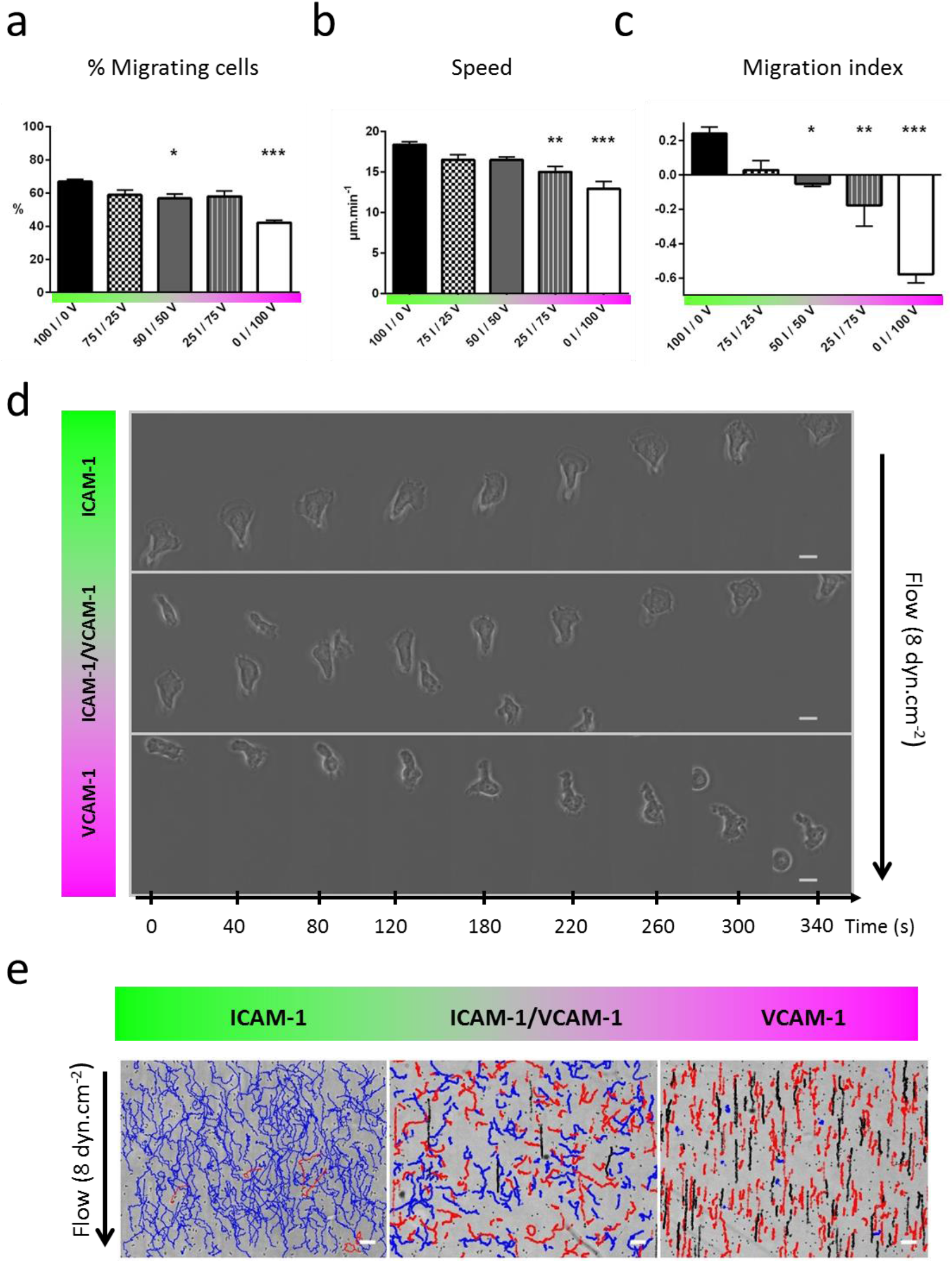
Flow reveals different migration modes on mixed ICAM-1/VCAM-1 substrates. **a)** Percentage of migrating cells and **b)** Speed versus substrates composition in shear free conditions. x I / y V stands for X % ICAM-1 / y % VCAM-1. **c)** Population-averaged migration index (> 0 against the flow, <0 with the flow) versus substrates composition at shear stresses of 4 dyn.cm^-2^. All data are mean + s.e.m, n= 6 independent experiments with at least 500 cells in each experiment, *P < 0.05, **P < 0.001, ***P < 0.001 with respect to substrate composition, 100 % ICAM-1 /0 % VCAM-1, one way ANOVA with post hoc Dunnett’s test. d) Bright-field images sequence of cells crawling on pure ICAM-1, pure VCAM-1 and mixed ICAM-1/VCAM-1 substrates. Scale bar 10 μm, time laps 40 s. e) Trajectories of mobile cells on pure ICAM-1, pure VCAM-1 and mixed ICAM-1/VCAM-1 substrates, with a color code for cells crawling upstream (blue), crawling downstream (red) and rolling (black). Time span 17 mn, scale bar 100 μm.

### ICAM-1 imposes strong adhesion and crawling whereas VCAM-1 allows transient adhesion and rolling

Interactions with substrates coated by integrin ligands have little influence on motility machinery but a critical role on mechanotaxis response. To better understand the coupling between integrins and flow mechanotaxis, we further characterized adhesion properties. In absence of flow, the fraction of non-adherent cells decreased noticeably, and the fraction of immobile cells increased with the fraction of VCAM-1 on the substrates (Figure 2a,b). This apparent increase of adhesivity with VCAM-1 can mainly be attributed to a population of round cells. Round cells, as opposed to polarized cells, are not able to crawl and carry mostly low affinity integrins whose binding epitopes are hidden for LFA-1 and partially accessible for VLA-4. Weak adhesion is therefore possible for round cells with VCAM-1 and not with ICAM-1, which is consistent with a larger number of adherent cells on VCAM-1 in shear free conditions. The additional amount of adherent and round cells with VCAM-1 explains also the larger fraction of immobile cells without flow and the large fraction of immobile cells that detached on VCAM-1 under flow (Figure 2c). Immobile round cells adhering via low-affinity VLA-4 have indeed a low adhesivity. Conversely, Figure 2c shows also that immobile cells on ICAM-1 did not detach under flow, which is in accord with LFA-1 adhesion occurring only in high affinity state and being reinforced by ligand engagement. Let us consider now the case of mobile and polarized cells. Figure 2c shows a stronger adhesion on ICAM-1-rich than on VCAM-1-rich substrates. This difference is consistent with a larger expression of LFA-1 than VLA-4 revealed by quantitative flow cytometry (Supplementary Information 2), however other parameters are also determinant for adhesion control such as affinity, avidity or clustering properties, which differ between integrins. Interestingly, mobile cells only crawl and never roll on ICAM-1 (Figure 2d), which is in line with a robust adhesion mediated by ICAM-1/LFA-1. In contrast, the fraction of rolling cells increases with the fraction of VCAM-1, up to a maximum of 25% on pure VCAM-1 substrates, which reveals weaker and potentially transient adhesion. Altogether, these results on adhesion strength of immobile and motile cells support that ICAM-1 promotes strong adhesion and a high propensity for crawling, whereas VCAM-1 promotes weaker and more transient adhesion, yielding a coexistence of rolling and crawling cells.

**Figure 2:**
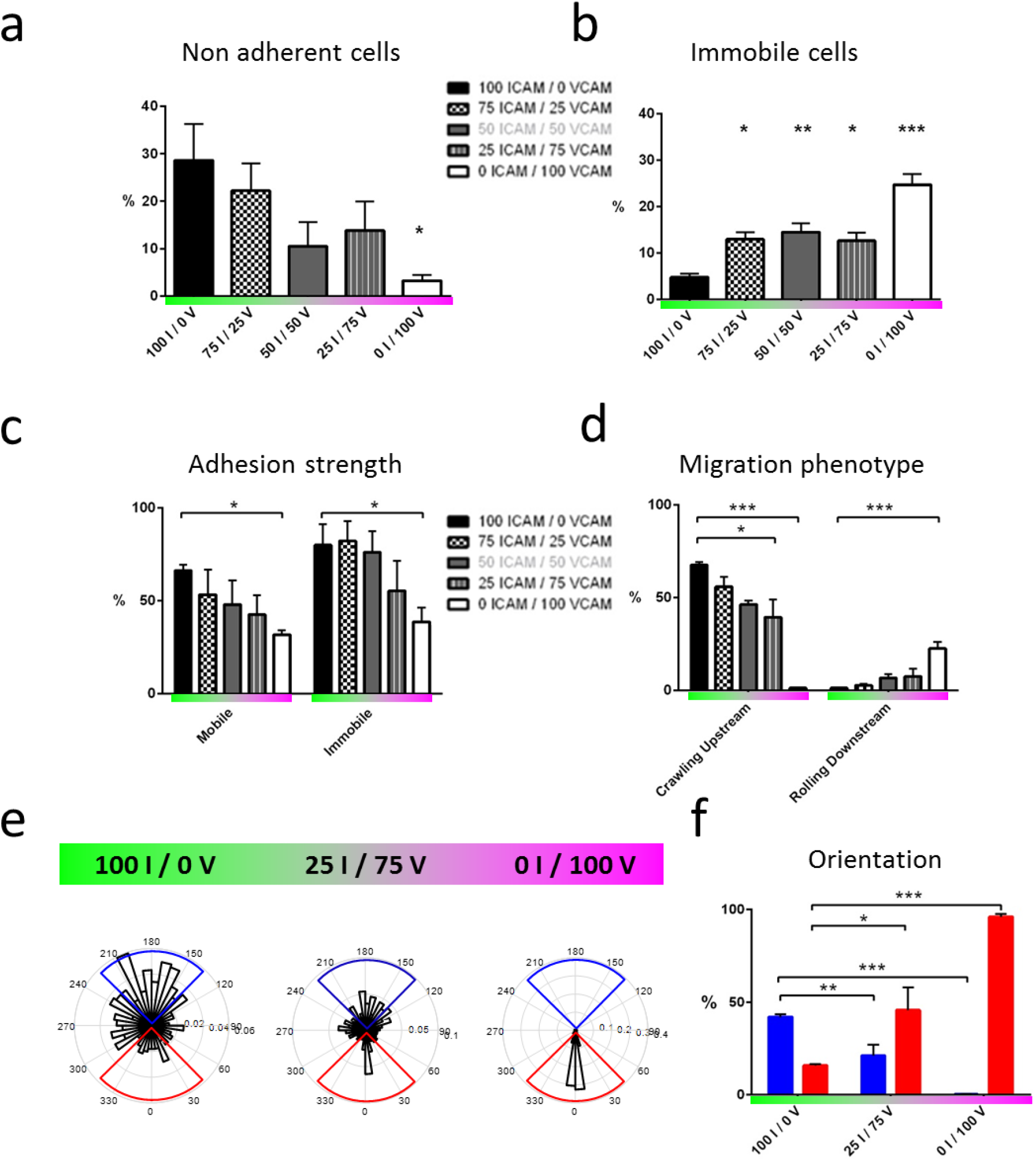
ICAM-1 imposes strong adhesion and upstream crawling whereas VCAM-1 allows transient adhesion and downstream crawling/rolling. **a)** Percentage of non-adherent cells and **b)** immobile cells versus substrate composition in shear free condition. **c)** Adhesion strength of mobile and immobile cells, measured as the percentage of cells resistant to a shear stress of 4 dyn.cm^-2^ with respect to the initial number of cells on the substrate. **d)** Percentages of cells crawling upstream and rolling downstream with respect to the total number of cells migrating on the surface, under a shear stress of 4 dyn.cm^-2^ and for different substrate composition. **e)** Rose plots of cells directions at different substrate composition. **f)** Percentage of upstream (blue) and downstream (red) crawling cells, determined here by cumulating data in the blue and red quadrant of the rose plots of cell migration for respectively upstream and downstream crawling cells. All data are mean + s.e.m, n= 6 independent experiments with at least 500 cells in each experiment, *P < 0.05, **P < 0.001, ***P < 0.001 with respect to substrate composition, 100 % ICAM-1 / 0 % VCAM-1, one way ANOVA with post hoc Dunnett’s test.

### ICAM-1 promotes upstream crawling and VCAM-1 allows downstream crawling

To further characterize the guiding of crawling cells versus integrin-ligand involved, we quantified the population of cells with downstream or upstream phenotype by taking into account trajectories making an angle ±45° respectively with and against the flow direction (Figure 2e,f). Upstream phenotype was maximum on pure ICAM-1 substrates (50 % at 4 dyn.cm^-2^) and absent on VCAM-1 substrates, which suggests that upstream phenotype is associated to ICAM-1-mediated cell adhesion. In contrast, the fraction of cells with downstream phenotype increased with the fraction of VCAM-1 on the substrates up to 100 % on pure VCAM-1 substrates, which suggests that VCAM-1 mediates downstream phenotype or inhibits upstream phenotype. These general trends are robust, however they do not explain the biphasic behavior on mixed substrates. A finer characterization of the properties of upstream and downstream phenotypes at the single cell level is necessary to shed light on opposite behaviors in a given population.

### Speed remains constant for upstream crawling cells

Figure 3a shows that the speed of rolling cells increased with shear stress on mixed ICAM-1/VCAM-1 substrates as well as on VCAM-1 substrates, which is trivial since rolling is passively powered by the flow push on transiently adherent cells. In contrast, the upstream crawling T cells on ICAM-1 substrates had a constant speed relative to the shear stress despite opposed action of flow. The apparent slight decrease of speed on ICAM-1 substrates was previously identified as a population selection effect, and the velocity of single cells was shown^12^ to be constant up to shear stress of 60 dynes.cm^-2^. The reason is that the hydrodynamic force on a whole cell (< 0.1 nN) is negligible as compared to the force developed by cells crawling machinery (several nN)^12^. Interestingly, cells crawling upstream on mixed ICAM-1/VCAM-1 substrates had also a constant velocity versus flow, exactly like on pure ICAM-1 substrates. Upstream crawling is therefore characterized by strong adhesion strength and migration power on both ICAM-1 and mixed ICAM-1/VCAM-1 substrates.

**Figure 3:**
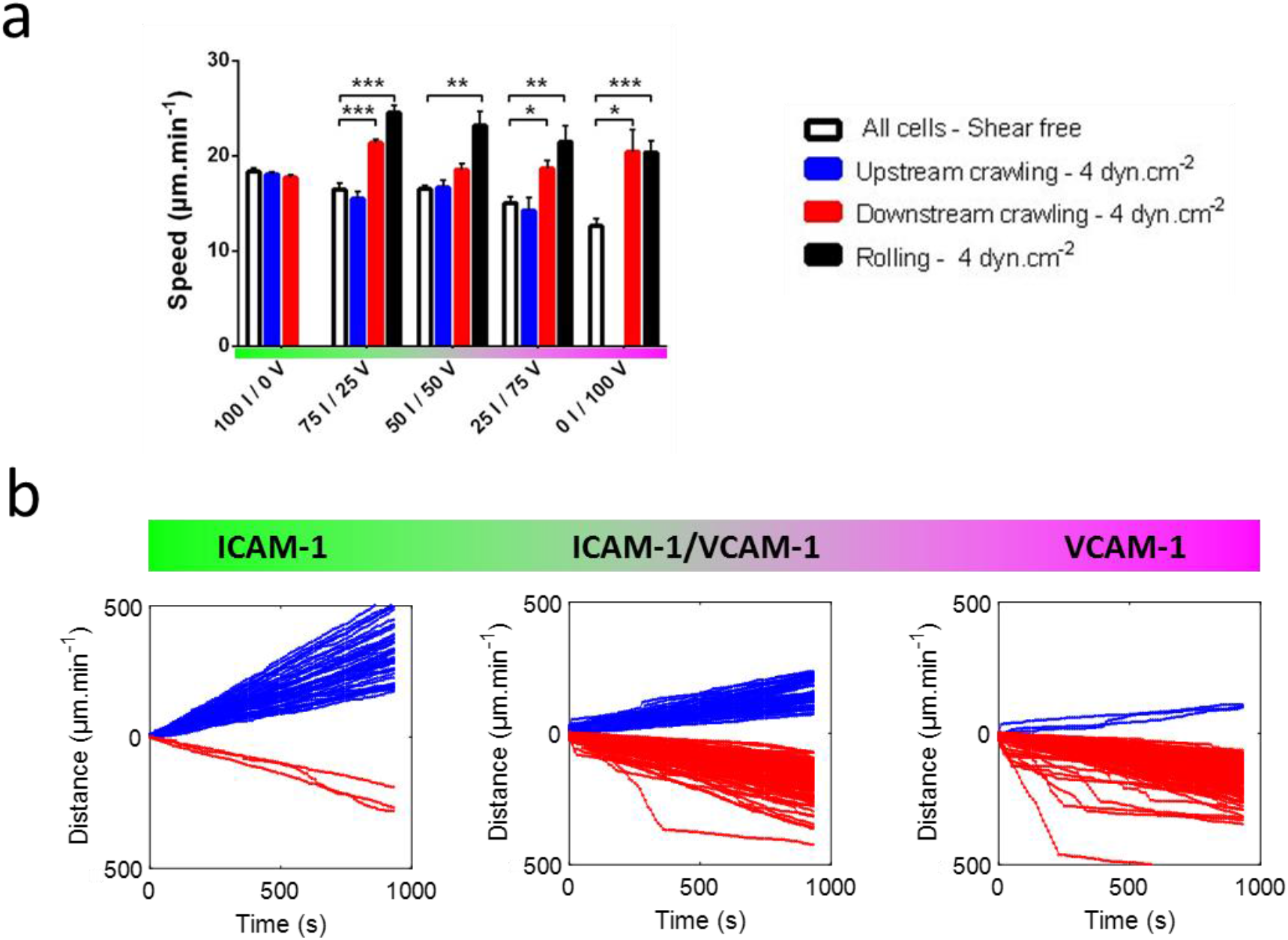
Speed increases with flow for rolling cells but remains constant for upstream crawling cells. **a)** Speed versus substrate composition of all cells in shear free condition, and of upstream crawling cells and downstream crawling cells and rolling cells under a shear stress of 4 dyn.cm^-2^. x I / y V stands for X % ICAM-1 / y % VCAM-1. All data are mean + s.e.m, n= 6 independent experiments with at least 500 cells in each experiment, *P < 0.05, **P < 0.001, ***P < 0.001, one way ANOVA with post hoc Dunnett’s test. **b)** Cumulative distance travelled by individual crawling cells on ICAM-1 (left), mixed ICAM-1/VCAM-1 (center), and VCAM-1 (right) substrates. The color of each curve indicates the migration mode of the corresponding cell tracked, blue for upstream and red for downstream crawling cells.

### Speed increases with flow for downstream crawling cells

For cells crawling downstream, the population-averaged data (Figure 3a) shows a significant increase of speed for downstream crawling when flow increases. Since flow actuation is orders of magnitude weaker than the power of the crawling machinery, this effect is not a straightforward action of flow push. A closer look at individual cell tracks (Figure 3b) revealed that curvilinear displacements vs. time were linear (constant slope) for cells crawling upstream (on ICAM-1 and mixed ICAM-1/VCAM-1), and composed of segments with different slopes for cells crawling downstream (on VCAM-1 and mixed ICAM-1/VCAM-1). Hence, in the case of upstream crawling cells the speed was different from one cell to another but constant vs. time for a given cells. This observation is in line with a stable motility machinery and a negligible effect of flow on cell crawling. In the case of downstream crawling, slow sequences corresponded to crawling (Supplementary Movie 3), with the slower motion as compared to upstream crawling being attributed to a higher tendency of cell rear attachment, whereas the fast sequences correspond to mixed crawling and rolling by detachment of cell rear. This observation is consistent with VLA-4 mediating transient and less robust adhesion than LFA-1.

### Uropod is detached for upstream bound cells and attached for downstream-bound cells

To shed further light on the mechanism underlying orientation under flow, we used Reflection Interference Contrast Microscopy (RICM) to image the adhesion footprint of cells during migration. Uropods were found non-adherent for upstream crawling cells and markedly adherent for downstream crawling cells for all types of substrates coatings (Figure 4 and Supplementary Movie 4). These observations support that the uropod-wind vane mechanism functions independently of the substrate composition. Cells migrate upstream whenever the uropod-wind vane mechanism is ON (detached uropod). Conversely, they have no reason to go upstream when the uropod-wind vane mechanism is OFF (attached uropod). An alternative mechanism must however be at work to foster the orientation of downstream crawling cells. Figure 4 shows that lamellipods, which were strongly adherent for cells crawling upstream, were markedly non-adherent for cells crawling downstream. As previously suggested for neutrophils^11^, a lamellipod loosely connected to a substrate is passively funneled by flow, yielding a preferential downstream orientation to the cell body. Altogether, the characteristics of cell adhesion patterns on ICAM-1 and VCAM-1 substrates indicate that two mechanisms are at work to guide cells versus flow: the uropod-wind vane mechanism guides cells upstream whenever the uropod is detached, and a lamellipod focusing mechanism guides cells downstream whenever the uropod is attached. These two mechanisms based on cell adhesion footprint provide a self-consistent mechanistic picture for upstream and downstream mechanotaxis phenotypes. Noticeably, both mechanisms are passive in the sense that they require a priori no mechanotransduction for re-orientation, but merely a mechanical orientation of the cells parts that are loosely attached.

**Figure 4:**
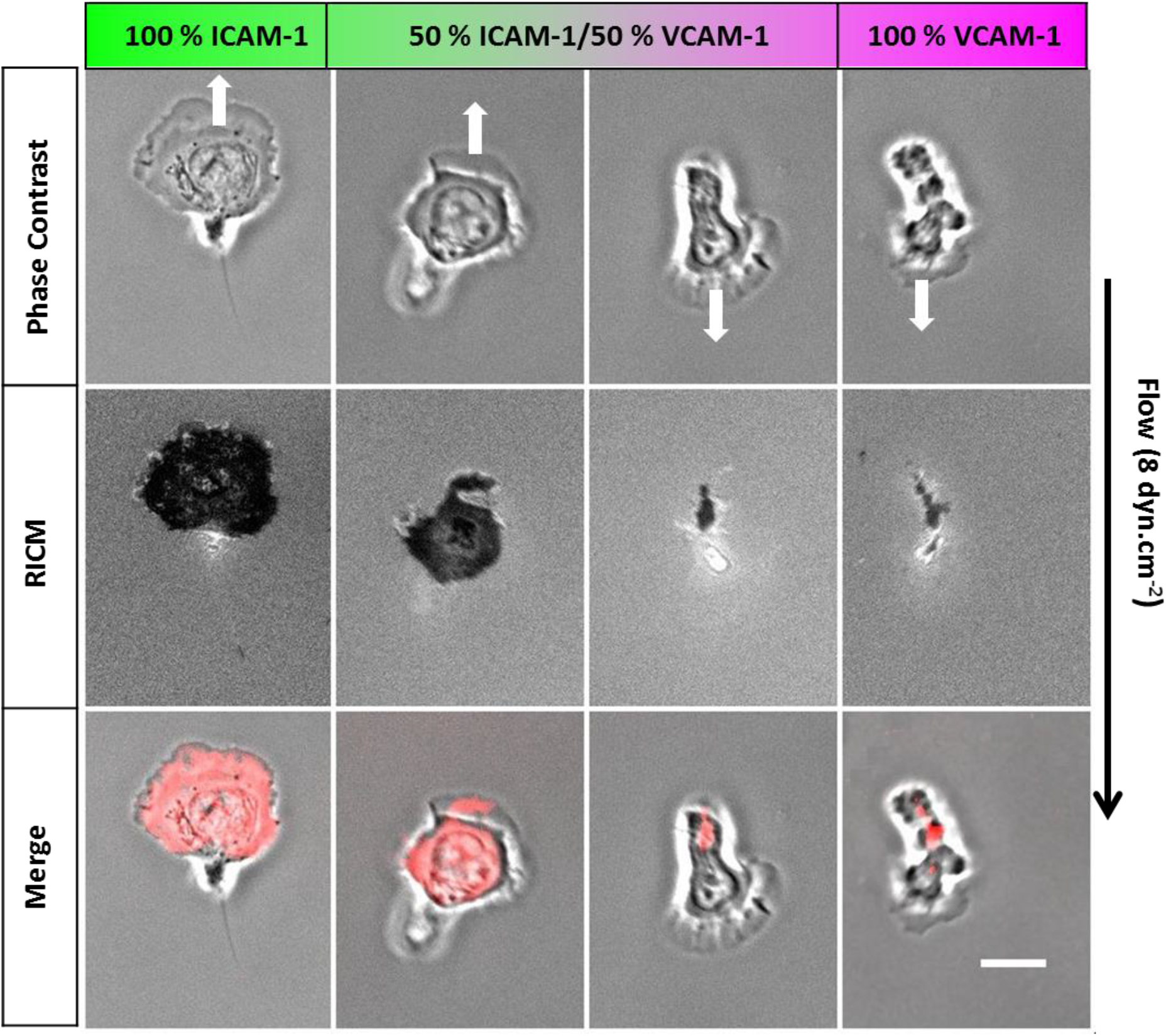
Cell rear is detached for upstream bound cells and cell front for downstream-bound cells. Image sequences of crawling cells under a flow of 8 dyn.cm^-2^ in phase contrast (top), reflection interference contrast RICM (center). On the merged images, the RICM image is contrast-inverted and colored in red. The black arrow indicates flow direction and white arrow direction of cell migration. The adhesion zone (dark in RICM, red in merge) is positioned in cell front for upstream crawling cells and in cell rear for downstream crawling cells. Scale bar 10 μm.

### Flow triggers no calcium signaling

Although passive mechanisms explain cells orientation under flow, one cannot discard a role of active mechanisms based on signaling triggered by integrins or other mechanotransduction events. We therefore monitored the intracellular calcium activity during flow stimulation. Figure 5 (and Supplementary Movie 5) shows that calcium activity upon onset of flow remained below detection level on all substrates coated with ICAM-1 and/or VCAM-1, whereas control with ionomycin showed a strong signal. Since calcium signaling is shared by many intracellular signaling pathways^35^, these data support that mechanotransduction is not involved in T cells guidance by flow.

**Figure 5:**
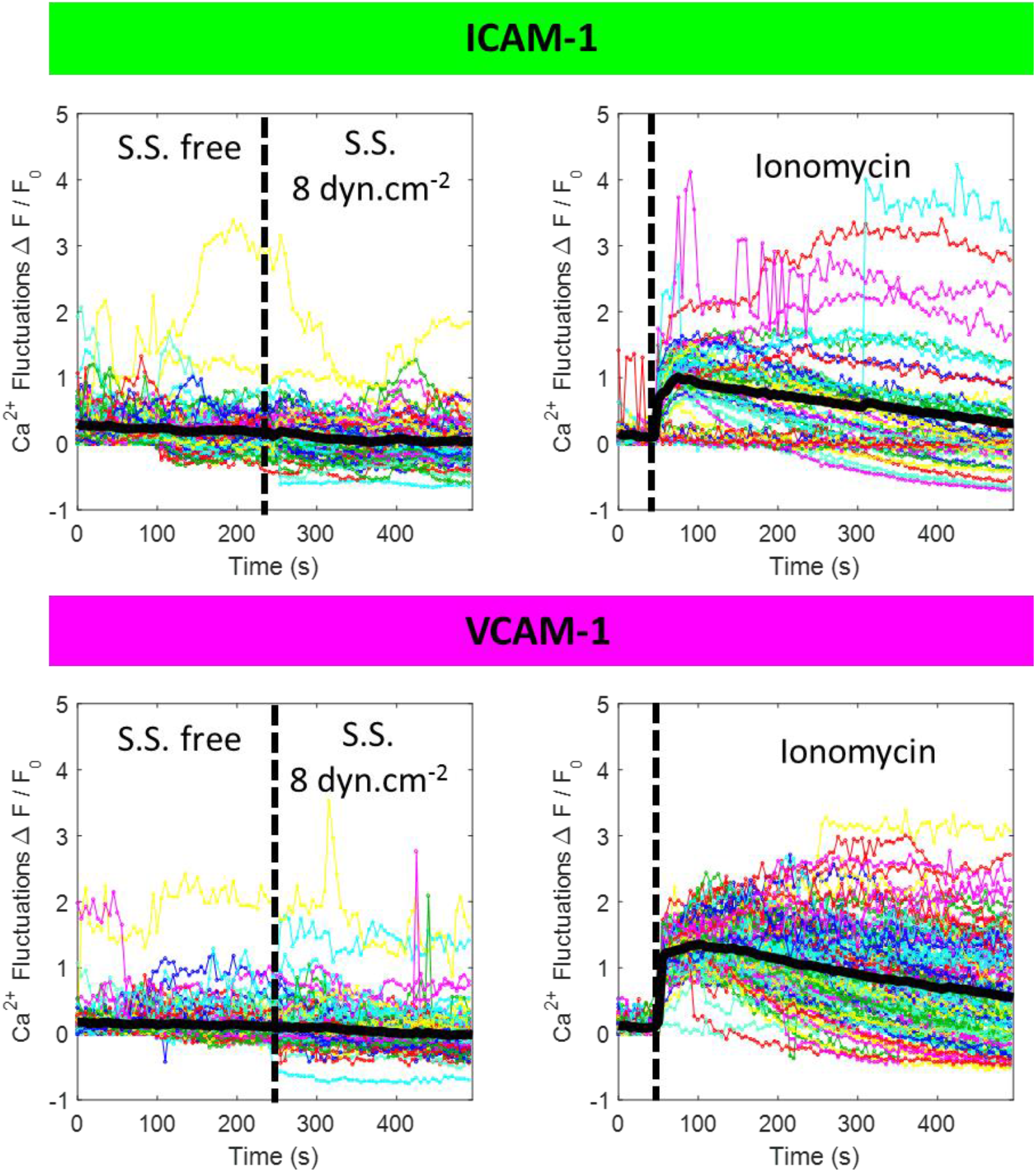
Absence of calcium signaling triggered by flow supports absence of mechanotransduction in flow mechanotaxis. Calcium signaling versus onset of flow (left) or addition of ionomycin (right) in cells loaded with Oregon Green 488 BAPTA-1 and crawling on ICAM-1–top) or VCAM-1 (bottom) substrates.

### The affinities of LFA-1 and VLA-4 are polarized in opposite direction

RICM revealed different positioning of the cell adhesion depending on substrate types, but it provided no information on the type and the amount of integrins involved in the adhesion zones. To perform functional imaging of adhesion footprints under flow, we used Total Internal Reflection Fluorescence microscopy (TIRF) and the antibodies mAb 24 and mAb B44 against respectively integrins LFA-1 and VLA-4 in high affinity states (Figure 6a). TIRF experiments had to be performed on fixed cells because antibodies altered strongly and instantly the regulation of integrins affinity in live conditions. Adhesion of frontward zone on pure ICAM-1 contained mostly high affinity LFA-1, and adhesion of rearward zone on pure VCAM-1 mostly high affinity VLA-4. The absence of high affinity LFA-1 in VCAM-1-mediated contact zone and of high affinity VLA-4 in ICAM-1-mediated contact zones suggests that activation of integrins in contact zones requires local engagement with their respective ligand. However, ligand-induced activation does not promote adhesion of cell rear on ICAM-1 substrates nor of cell front on VCAM-1 substrates. Ligand induced activation is therefore not the sole mechanism involved. There must be up-regulation of the affinity of LFA-1 in cell front and of VLA-4 in cell rear, and conversely down-regulation of the affinity of LFA-1 in cell rear and of VLA-4 in cell front. In any case, LFA-1 and VLA-4 have their affinity strongly polarized from cell front and rear, and with opposite direction.

**Figure 6:**
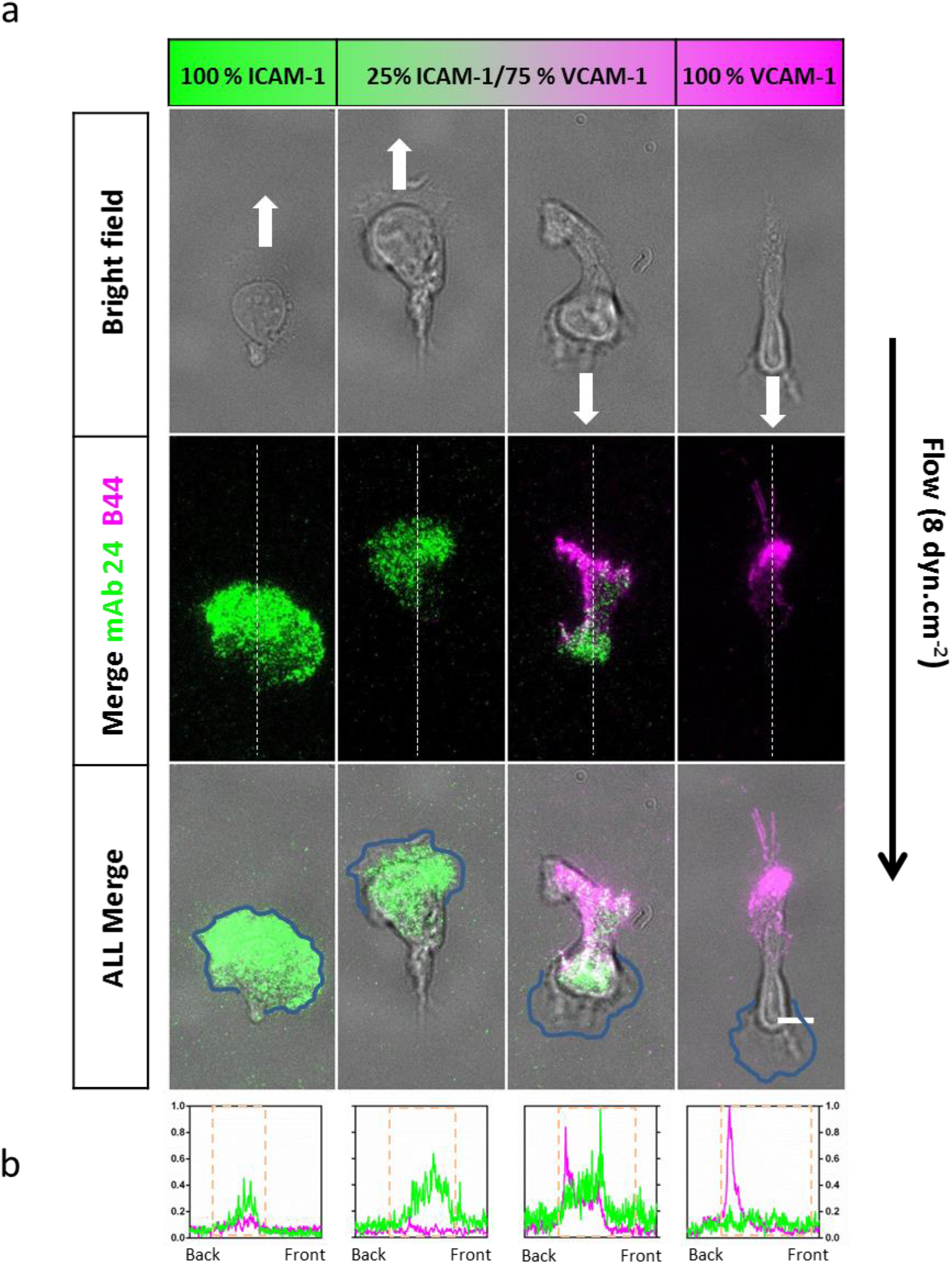
Imaging of high affinity LFA-1 and VLA-4 in cell contact zone by TIRF reveals complex regulation mechanism of their affinity. **a)** Microscopic images in bright field (top) and TIRF (center) of crawling T cells fixed under a shear stress of 8 dyn.cm^-2^ and stained for high affinity LFA-1 with mAb24 (green) and high affinity VLA-4 with mAb B44 (magenta). Scale bar 10 μm. **b)** Histograms performed for each fluorescent channel, to highlight integrin distribution along the cell axis. Values were normalized to the highest value recorded on either condition. Dashed boxes indicate the cell body area.

### Bistability between upstream crawling with LFA-1 and downstream crawling with VLA-4

On mixed ICAM-1/VCAM-1 substrates, both integrins are stimulated by their ligand in the contact zones, so that cells with high affinity LFA-1 in their front and high affinity VLA-4 in their rear could a priori adhere by their two poles. Considering upstream crawling cells, RICM imaging showed in fact that cells adhered only by their front, and TIRF revealed that such frontal adhesion zones involved almost exclusively LFA-1 (Figure 6b). Down-regulation of VLA-4 affinity by cell polarization signaling, previously evidenced, explains the absence of activated VLA-4 in cell front but not the absence of adhesion of cell rear. An additional mechanism is necessary to hamper the adhesion of cell rear by VLA-4 and an inhibiting crosstalk of activated LFA-1 towards VLA-4, already reported in the literature^222222222222^ can exactly play this role. Considering now downstream crawling cells, RICM showed that they are strongly attached by their rear. TIRF further revealed that high affinity VLA-4 was dominant in this posterior adhesion, whereas high affinity LFA-1 was mostly present in cell central zone and partially also in cell rear (Figure 6b). These observations go against an inhibiting crosstalk of activated VLA-4 toward LFA-1 and rather in favor of an activating crosstalk since LFA-1 affinity is usually downregulated in cell rear on ICAM-1 substrate. Altogether, these data show that polarization of integrins and ligand activation alone cannot explain differential orientation under flow and that other mechanisms involving crosstalk mechanisms between integrins are also involved.

### The level of high affinity integrins dictates orientation decision

Integrins LFA-1 and VLA-4 are crucial in cell orientation under flow, but the decision process to choose upstream or downstream orientation for a given cell remains unclear. To challenge the existence of populations with different LFA-1 and VLA-4 expression levels, we performed flow cytometry experiments with double staining of αL (for LFA-1) and β2 (for VLA-4), and found a single population (Figure 7a). Opposite orientations under flow can therefore not arise from distinct populations with sharply different levels of integrins LFA-1 and VLA-4 expression. However, a bistable system can trigger sharply distinct responses within a single cell population with distributed LFA-1 and VLA-4 expressions. To challenge this mechanism, we performed perturbation experiments to address the correlation between effective integrins levels and migration phenotype under flow. The effective number of available LFA-1 and VLA-4 integrins at cell surfaces was tuned by addition of blocking antibodies against high affinity LFA-1 or VLA-4 (Figure 7b). On mixed ICAM-1/VCAM-1 substrates, the blocking of 50 % of LFA-1 integrins displaced the phenotypes distribution from upstream to downstream migration, and conversely the blocking of 50 % of VLA-4 integrins displaced the phenotypes distribution from downstream to upstream migration (Figure 7c and Supplementary Movie 6). These data support directly the hypothesis that the bistable choice of a given cell to go up- or down-stream depends on its relative amounts of VLA-4 and LFA-1. A cell rich in LFA-1 will adopt the upstream migration phenotype like T-cells on ICAM-1 substrate, and a cell rich in VLA-4 will adopt the general downstream migration phenotype like T-cells on VCAM-1 substrates.

**Figure 7:**
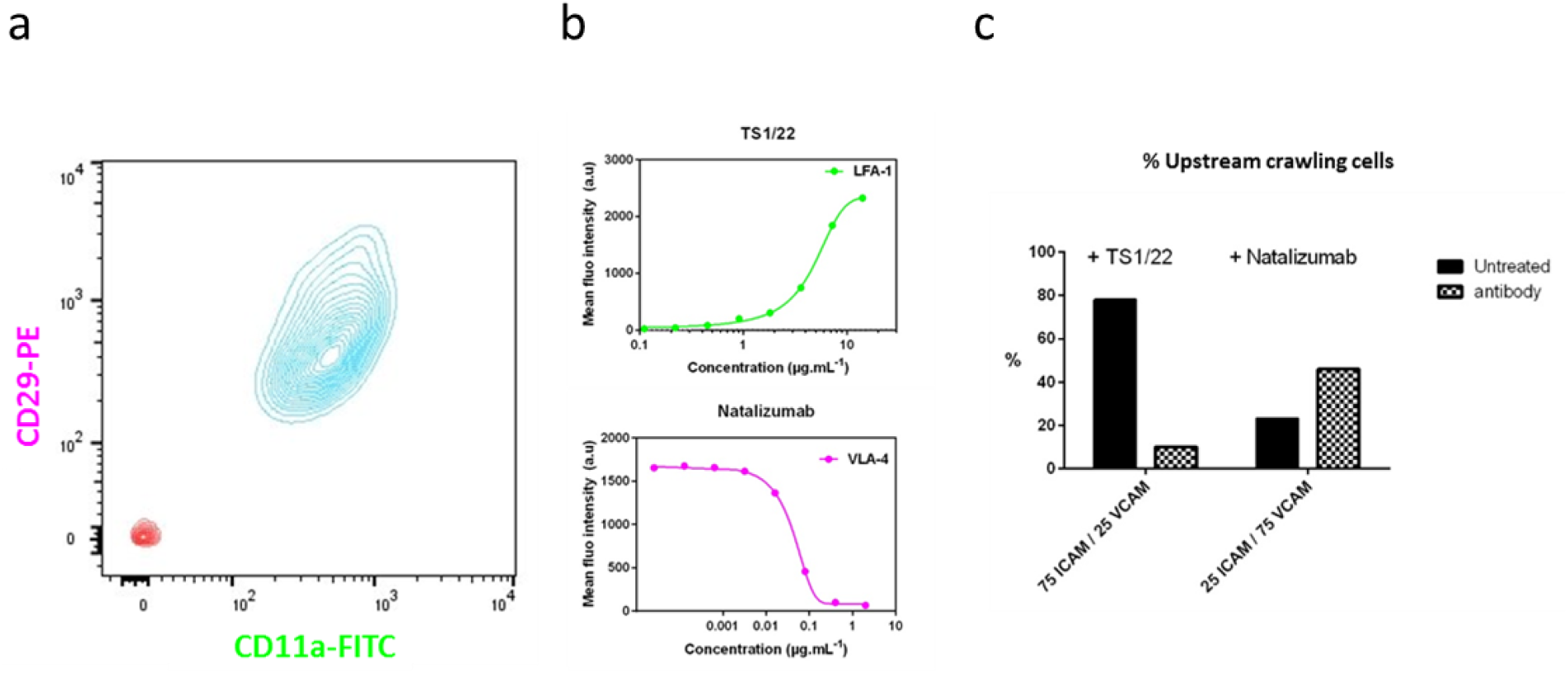
Perturbation experiments support that integrin expression level dictates sharp decision of orientation versus flow. **a)** 2D cytometry graphs of activated T cells versus expression of heterodimer αL (Ab a-CD11a) for LFA-1 and β2 (Ab α-CD29) for VLA-4. **b)** Cytometry data of blocking antibody TS1/22 against LFA-1 and Natalizumab against VLA-4 linked to effector T lymphocytes versus concentration. **c)** Percentages of upstream crawling cells on mixed ICAM-1/VCAM-1 substrates with and without addition of blocking antibodies TS1/22, against integrins LFA-1 (left), and Natalizumab, against VLA-4 (right). Blocking of LFA-1 displaces phenotype distribution towards downstream phenotype, and blocking of VLA-4 towards upstream phenotype.

## DISCUSSION

### Bistability generates a biphasic system for crawling T cell under flows

Upon recruitment from the blood stream, lymphocytes crawl on the intraluminal surface of blood vessels presenting noticeably ICAM-1 and VCAM-1 adhesion molecules under a shear stress of 5-10 dynes.cm^-2^ ^36^. Shear stress has recently been recognized in vivo and in vitro as an efficient stimulus to guide crawling T lymphocytes, albeit function and mechanisms of these guiding phenomena remain poorly understood. We confirmed here that lymphocytes display upstream crawling on ICAM-1 ^10,12–14,37^, downstream migration by rolling and crawling on VCAM-1 ^14,26^, and that they change from upstream to downstream migration when the ratio of VCAM-1 over ICAM-1 increase^14^ on average population level. Our analysis at single cell level has however shown that individual cells do not adopt intermediate phenotypes between upstream and downstream but choose either one of these opposite phenotypes. Instead of a smooth transition of phenotype by a homogenous population, we revealed a biphasic system with two distinct populations of upstream and downstream phenotypes. From a physical point of view, the separation of a system in two distinct phases may rely on a first order phase transition or on a bistable mechanism. There is no thermodynamic-like phase transition here because individual cells do not exchange between the two different states (phenotypes). A bistable mechanism is however plausible because individual cells ends up and remain in distinct states. We deciphered indeed the chain of mechanistic elements at the molecular and cellular level that allow the emergence of a bistable system.

### A precise cartography of integrins affinity is delineated

Integrins regulation plays a central role in cells response to flow and our results bring new insight in the spatial distribution of high affinity integrins along polarized migrating T cells. Previous investigations on ICAM-1 substrates had reported a spatial segregation of LFA-1 affinity state, with lamellipod corresponding exclusively to intermediate states and high affinity states confined in the central or “focal” zone^38^. In this work, we find that high affinity LFA-1 are indeed strongly present in the focal zone but also the lamellipod of T cells on ICAM-1 substrates, and more generally for upstream-bound T cells. In contrast, we also evidenced that high affinity LFA-1 are localized in the central and rear zones of downstream bound cells. This more precise cartography of integrins affinity may result partly form the improvement of staining protocols with mAB 24 but also from the distinctive analysis of sub-populations of T cells with different phenotypes of integrin-mediated mechanotaxis.

### T cells orient under stress without mechanotransduction

Guiding of migrating cells by an external stimulus recalls a sophisticated mechanism evolved for a specific function. Chemotaxis involves stimuli detection and signal transduction by molecular receptors followed by internal signal treatment to reorganize the cytoskeleton and trigger cell orientation^1^. For mechanotaxis, several observations supports that similar complex mechanisms involving molecular mechanodetection and mechanotransduction allow deterministic cells guiding. For adherent endothelial cells, shear stress–induced orientation involves signaling via integrins subunits α5 and β1^34^. Molecular basis of mechano-signaling via integrins and further downstream signaling events have also been identified, like the dephosphorylation of p130CAS under tension, and inactivation of Rho-GTPase Rac1 in the side of cell facing flow^33^. It was further demonstrated *in vitro* that CAS substrate domain acts as a primary molecular force sensor, transducing force into mechanical extension and thereby priming phosphorylation and activation of downstream signaling^32^. Mechanotransduction by integrin adhesion complexes seem in the end to play a central role in mechanotaxis of cells forming focal adhesion complexes. In contrast, amoeboid cells like leukocytes migrate at high velocities without maturation of integrin-mediated focal contacts into focal adhesions. The role of mechanotransduction by integrins remain therefore more arguable. For downstream mechanotaxis of neutrophils, Dixit et al ^21^ hypothesized that shear forces used high-affinity LFA-1 transmission to facilitate the cooperation with the calcium release-activated channel Orai1 in directing localized cytoskeletal activation and subsequent directed migration. Besides, Artemenko et al^35^ or NIetmhammer^35,39^ have shown that flow could activate internal signalling networks common with chemotaxis. Several elements are therefore supporting the hypothesis that leukocyte flow guiding may be mediated by active signal transduction and processing, like for chemotaxis. In contrast, we previously proposed a model without mechanotransduction for upstream migrating effector T lymphocytes under flow mediated by one integrin LFA-1^13^. This alternative mechanism was based on two established properties of crawling effector T cells, a detached tail (or uropod) acting as a wind vane, and a robustly maintained front-rear polarization. These two elements operate also in the absence of flow and this mechanotaxis mechanism requires no signal triggered by flow. This model was extended here to a more complex system of flow mechanotaxis controlled by two integrins, LFA-1 and VLA-4, and with a choice between two directions. The mechanism of uropod/wind-vane was still found operative whenever the cell uropod was detached, promoting cells migration against the flow. A complementary mechanism of lamellipod flow focusing was shown to interfere with loosely adherent lamellipod tip, and to promote cells migration with the flow, as already observed with keratocytes^40^. Altogether, upstream and downstream mechanotaxis of effector T lymphocytes adhering via LFA-1 or VLA-4 integrins can rely on passive mechanisms without mechanotransduction.

### Bistability relies on opposite spatial regulation of integrins affinity

At the cellular level, the mechanisms of uropod/wind vane and lamellipod flow focusing explain upstream and downstream phenotypes for all cells and all compositions of ICAM-1/VCAM-1 substrates, but not how a phenotype and its associated mechanism are chosen by individual cells. The explanation was found at the molecular level. Bistability of cell orientation results from bistability of cell adhesion footprint in front or rear pole, which relies itself on the oppposite polarization of LFA-1 and VLA-4 affinities along the rear-front axis of cells. High affinity polarization was found to point forward for LFA-1 and backwards for VLA-4, which agrees with literature data reporting affinity upregulation for LFA-1 in cell front^38,41^ and of VLA-4 in cell rear ^42^ on one hand, and affinity downregulation of LFA-1 in cell rear^38,43,44^ and of VLA-4 in the cell front^22^ on the other hand. These opposite polarizations of integrins explain directly the selective anterior adhesion of cells on ICAM-1 with associated upstream phenotype, and the selective posterior adhesion of cells on VCAM-1 with associated downstream phenotype. On mixed substrates, perturbation experiments with blocking antibodies of LFA-1 and VLA-4 allowed us to establish a link between the cell orientation under flow and the implication of LFA-1 or VLA-4 in cell adhesion. Higher levels of high affinity LFA-1 vs. VLA-4 on cells and/or of higher ratio ICAM-1 vs. VAM-1 on substrates favored the upstream state, whereas lower expression of LFA-1 vs. VLA-4 and/or lower ratio of ICAM-1 vs. VAM-1 on substrate favors the downstream state (Figure 8).

**Figure 8:**
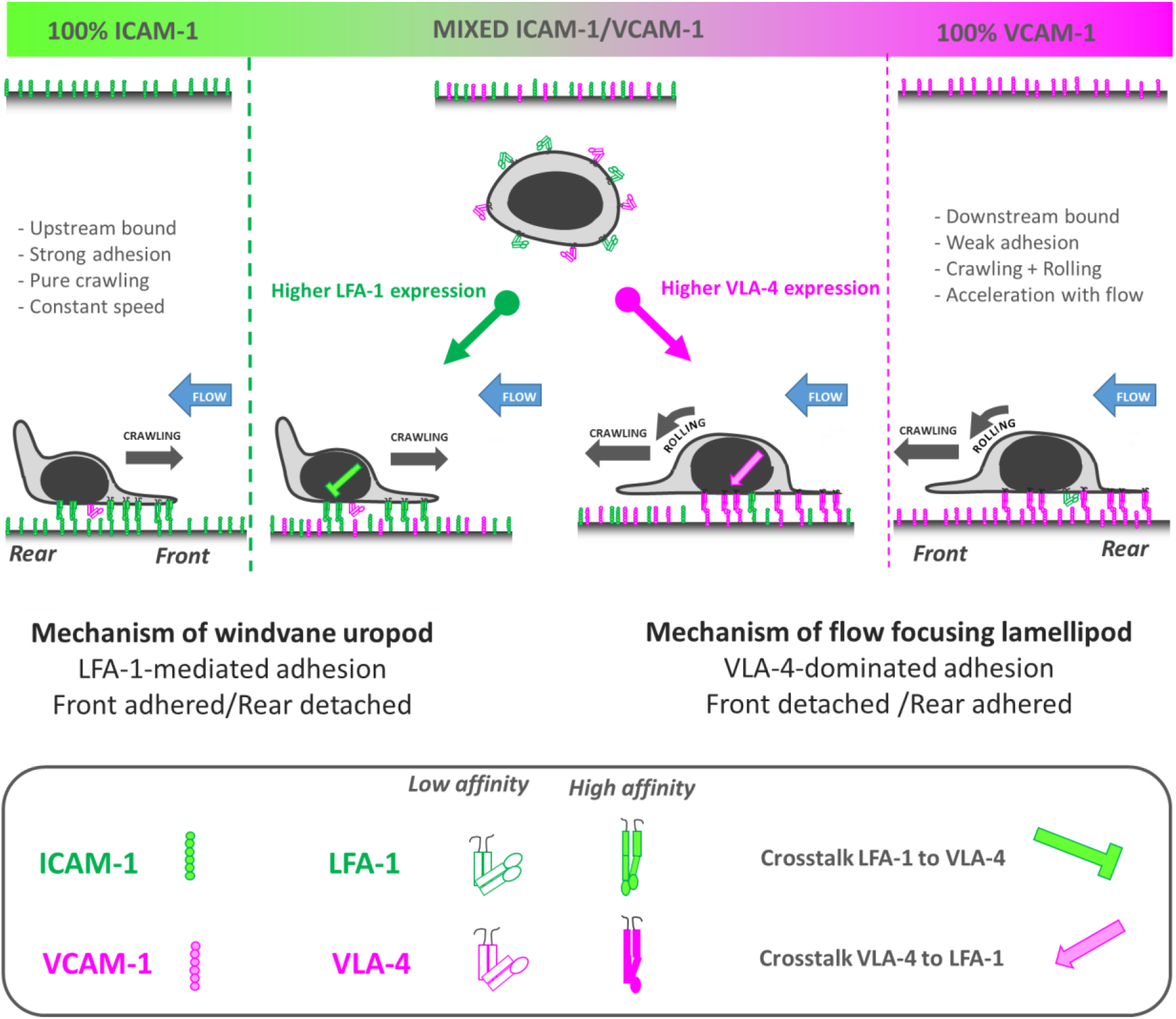
A bistable mechanism of cell adhesion spatial regulation explains integrin control of T cell flow mechanotaxis. On pure substrates of ICAM-1 or VCAM-1, T cells population have a homogeneous phenotypes with opposite orientation on ICAM-1 and VCAM-1. On mixed substrates of ICAM-1 or VCAM-1, T cells distribute in two populations with opposite orientations and characteristics similar to phenotypes on pure substrates. Decisions of orientation on mixed substrates are controlled by the expression level of integrins LFA-1 and VLA-4 via a bistable polarization of cell adhesion: a higher LFA-1 expression leads to a LFA-1-dominated adhesion of cell front (very similar to upstream crawling cells on ICAM-1), whereas a higher expression of VLA-4 leads to adhesion of cell rear and center (very similar to downstream crawling cells on VCAM-1). Inhibiting crosstalk of LFA-1 towards VLA-4 reinforces adhesion polarization toward cell front, which favors wind vane mechanism and upstream phenotype. Activating crosstalk of VLA-4 towards LFA-1 reinforces adhesion of cell uropod, which hampers wind vane mechanism and favors downstream phenotype.

### Bistability requires integrins crosstalk

Attachment of a single cell pole is essential to explain system bistability but polarization of integrins affinity cannot sustain it alone. Attachment of cells only by their front via LFA-1 on mixed substrates, revealed in our experiments for upstream cells, requires an inhibitory crosstalk of activated LFA-1 toward VLA-4 to detach the uropod. This crosstalk, which plays a crucial role in our mechanistic model of upstream guiding, has been evidenced in the literature^22,45^. It acts as an amplifier of integrin affinity imbalance, allowing cells with dominant LFA-1 vs. VLA-4 to behave like cells bearing only LFA-1 4 (Figure 8). For the case of downstream crawling cells, a symmetric inhibitory crosstalk effect from VLA-4 toward LFA-1 could promote symmetric orientation versus flow for cells with a dominant VLA-4 vs. LFA-1. However, we observed the presence of high affinity LFA-1 and VLA-4 in cell rear on mixed ICAM-1/VCAM-1. Since Polarization signaling inhibits LFA-1 in cell rear, our work suggests an activation crosstalk of VLA-4 toward LFA-1 that counterbalances polarization effect. This conclusion is also consistent with literature data showing that VLA-4 promotes activation of LFA-1^46,47^. Attachment of cell rear is therefore reinforced by the combined adhesion of VLA-4 and LFA-1, which inhibits totally wind vane mechanism and therefore the upstream phenotype. Nevertheless, provided that cell rear is attached, the extreme front edge of lamellipod, before its attachment, can promote downstream guiding. Figure 8 summarizes the interplay between integrins affinity regulation and cell directed migration choices under flow, and illustrates a unique model linking consistently mechanisms from the molecular to the cellular levels. Flow mechanotaxis decisions and bistability of guidance under flow rely on a polarized inside-out regulation of integrins LFA-1 and VLA-4 affinity and on an inhibitory crosstalk mechanism between LFA-1 and VLA-4 at the molecular levels, relayed by a wind vane uropod mechanism and a lamellipod flow focusing at the cellular level. Altogether, several integrins working synergistically can mediate multiphasic mechanotaxis as switchable immobilizing anchors and not as force transducers^50^.

### Flow mechanotaxis is another function controlled by integrins in leukocyte recruitment

During recruitment of leukocytes from the blood system to inflamed zones, integrins are known to control several crucial functions. First for cell arrest in blood vessels, VLA-4 (α_4_β_1_) contributes to rolling and LFA-1 (α_L_β_2_) is essential for crawling^19,36^. Then for cell extravasation, endothelial overexpression of integrins ligands is arguably guiding leukocytes into specialized portals of transmigration^48^. Finally for tissue migration, integrins α_V_ condition the proper homing of lymphocytes to inflamed zones^48^. The mechanism of lymphocyte guidance by flow enriches therefore the panel of integrins functionalities in the sequence of leukocyte recruitment. Although passive mechanisms suggest the possibility of a fortuitous phenotype^13^, the robustness and sophistication of a mechanism with synergistic regulation and crosstalk of multiple integrins supports instead a system evolved for a given function in leukocyte recruitment. A complete knowledge of integrins functions in leukocyte recruitment is also valuable for therapeutic purposes, to modulate the immune response like in the treatment of multiple sclerosis by integrins blocking antibodies^49^.

## MATERIAL AND METHODS

### Cell culture

Whole blood from healthy adult donors was obtained from the “Établissement Français du Sang”. Peripheral blood mononuclear cells (PBMCs) were recovered from the interface of a Ficoll gradient (Eurobio, Evry, France). T cells were isolated from PBMCs with Pan T cell isolation kit (Miltenyi Biotec, Bergisch Gladbach, Germany), then were stimulated for 48 h with anti-CD3/anti-CD28 Dynabeads (Gibco by Thermo Fischer Scientific, Waltham, MA) according to the manufacturer’s instructions. T lymphocytes were subsequently cultivated in Roswell Park Memorial Institute Medium (RPMI) 1640 supplemented with 25 mM GlutaMax (Gibco, Gibco by Thermo Fischer Scientific, Waltham, MA, USA), 10% fetal calf serum (FCS; Gibco by Thermo Fischer Scientific, Waltham, MA, USA) at 37°C, 5% CO_2_ in the presence of IL-2 (50 ng.mL^-1^; Miltenyi Biotec, Bergisch Gladbach, Germany) and used 6 to 10 days after stimulation. At the time of use, the cells were >99% positive for pan-T lymphocyte marker CD3 and assessed for activation and proliferation with CD25, CD45RO, CD45RA and CD69 markers as judged by flow cytometry.

### Microscopy

Bright field, RICM and fluorescent images were performed on a Zeiss Observer Z1 microscope (Carl Zeiss, Oberkachen, Germany) piloted with μManager^49^. The microscope was equipped with a CoolSnap HQ CCD camera (Photometrics, Tucson, AZ, USA) and different Zeiss objectives (Plan-Apochromat × 10/0.3, × 20/0.8 and × 63/1.4 and NeoFluar 63/1.25 antiflex. The source was a CoolLED pE-300 (CoolLED, Andover, UK). A narrow band-pass filter (λ=546 nm ± 12 nm) was used for RICM.

TIRF images were recorded on Nikon Eclipse Ti microscope (Nikon Instruments, Europe), equipped with an ILas2 system (Roper Scientific, France) and controlled by Metamorph software (Molecular Devices, San José, CA). Images were taken with an Apo TIRF 60x/1.49 oil objective (Nikon), a Prime 95TM Scientific CMOS Camera (Photometrics, Tucson, AZ) and an Obis Coherent/ILAS LASER.

### Flow devices and surface treatment

Devices consisted in Ibidi μ-Slide IV^0.4^ uncoated (Ibidi GMBH, Martinsreid, Germany) and in homemade flow devices, fabricated using standard soft lithography routines^50^. Briefly, a positive mold was created with SU-8 2100-negative resins (Microchem, USA) on silicon wafers (Siltronix, France), then replicas were molded in polydimethylsiloxane (PDMS) elastomer (Sylgard 184, Dow Corning, USA) and sealed on glass coverslides via plasma activation (Harricks Plasma, Ithaca, NY, USA). Ports to plug inlet and outlet reservoirs were punched to a 1-mm outer diameter.

Flow devices (Ibidi μ-slide and homemade) were pre-coated for 1h at 37°C with 50 μg.mL^-1^ of Protein A (Sigma-Aldrich, St Louis, MO, USA). Surfaces were then blocked with 2.5% bovine serum album (BSA) (Axday, Dardilly, France) in PBS (Gibco, Gibco by Thermo Fischer Scientific, Waltham, MA, USA) for 30 min at room temperature. Channels were subsequently functionalized by an overnight incubation at 4 °C with 10 μg.mL^-1^ of either human ICAM-1-Fc or VCAM-1-Fc (R&D Systems, Minneapolis, MN, USA) in PBS or mixture of the CAMs. Channels were rinsed with PBS. Cells were then added in complete RPMI medium, allowed to equilibrate for 10 min and then rinsed with medium.

Cells in the flow chambers were observed at 37°C with the Zeiss Z1 automated microscope. Flow of pre-warmed and CO_2_ equilibrated culture media through the flow chamber was controlled using an Ibidi pump system (Ibidi GMBH, Martinsreid, Germany). Bright-field images (EC Plan-NEoFluar 10x/0.3 Ph1 objective) were collected every 10 s over the time frame indicated. The field of view represents 1740 × 1300 μm^2^.

### Fluorescence quantification of adhesion molecules

Anti-human CD106-PE and anti-human CD54-PE (eBioscience by Thermo Fischer Scientific, Waltham, MA) antibody were used for the quantification of substrates coatings with mixed ICAM-1 and VCAM-1. First we set up bulk calibration data by measuring the fluorescence intensity of 41 μm thick channels filled with antibody solutions at concentrations of 1.5, 3, 5 and 7 μg.mL^-1^. Channels were pretreated with 1% Pluronic F127 (Sigma-Aldrich, St Louis, MO, USA) fluorescence measured. Samples with mixed ICAM-1/VCAM-1 were then stained either with CD106-PE or with CD54-PE at 10 μg.mL^-1^ overnight at 4°C, and fluorescent images were then taken the next day and analyzed with Fiji software^51^. The average intensity at 5 different positions was converted into surface density using the bulk calibration data.

### Cell tracking and data analysis

A home-made program was developed with MATLAB software (The MathWorks, Natick, MA, USA) to track migrating cells and analyze their pathway properties. In all flow experiments, the flow is directed from the top to the bottom of the images presented here. To get an indication of the amount of motion in a particular direction, a migration index is calculated by dividing the distance the cells travel in the flow direction by the total distance travel by the cells. The average speed of a cell, *V*, is calculated as the ratio between the total trajectory length and the corresponding time of migration. All calculations were performed at least in triplicate for each substrate. Image Analysis was performed with ImageJ (U.S. National Institutes of Health, Bethesda, USA).

### Fixation under flow and Immunofluorescence staining

Cells migrating under flow were instantly fixed by quick injection of PFA 4% (Affymetrix, Cleveland, Ohio, USA) into the device. After 10 min incubation, the device was rinsed with PBS-Tween 0.1% and either stained or kept at 4°C until used. For the staining, cells were initially permeabilized with triton 0.5% (Sigma-Aldrich, St Louis, MO, USA) in PBS for 5 min and rinsed 3 times with PBS-Tween 0.1%. Free FC-binding sites of surface Protein A were blocked with human serum IgG 100 μg.mL^-1^ solution for 20 min; samples were then further blocked with BSA 2% for 20 min. After 3 washes with PBS-Tween, samples were incubated with mAb B44 (antibody at a concentration of 5 μg.mL^-1^ for 20 min, washed 3 times and then incubated with 2º anti-mouse (20 μg.mL^-1^) for another 20 min prior to imaging mAb B44 alone as a control. For the double staining of mAb B44 and mAb24, free binding sites of the secondary antibody were blocked by incubating the samples for at least 1hr with mouse IgG1 isotype control (10 μg.mL^-1^). Primarily labeled mAb24 antibody was finally added (4 μg.mL^-1^) and kept in solution during the imaging process.

The antibodies used were mouse anti-Integrin β1, clone name B44 (Millipore, Temecula, CA, USA); goat anti-mouse IgG, CF™647 conjugated (Sigma-Aldrich, St Louis, MO, USA); mouse IgG1 isotype control (Biolegend, San Diego, CA, USA); mouse anti-human CD11a/CD18, AF488 conjugated, clone name mAb24 (Biolegend, San Diego, CA, USA). All antibodies were diluted in PBS-Tween 0.1%. For the histograms of Figure 6-B, raw values were normalized by applying the following formula:

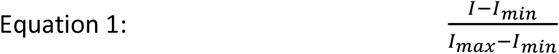

Where I_min_ is the minimal value recorded on each image and I_max_ is the maximum value recorded for each fluorophore in either condition.

### Fluorescent detection of Calcium Flux

For calcium imaging experiments, cells were first seeded in channels with RPMI medium and were incubated for 10 min at 37°C to allow adhesion, then they were rinsed with HBSS + 1% BSA and incubated with Oregon Green^®^ 488 BAPTA-1, AM (ThermoFisher, Waltham, USA) diluted in HBSS + 1% BSA (5 μM) for 15 min at 37°C in the dark. After rinsing with HBSS + 1% BSA, the medium was replaced by HBSS+ 10% SVF. Control experiment was achieved by injection ionomycin (ThermoFisher, Waltham, USA) at a concentration of 1 μg.mL^-1^.

### Flow Cytometry

One hundred thousand cells were taken from the cultured population and pelleted by centrifugation for 5 minutes at 1000 rpm. The cells were re-suspended in 100 μL PBS+2% FBS, containing the premixed antibodies (CD11a-FITC, clone Hi111 (eBioscience by Thermo Fischer Scientific, Waltham, MA, USA) and CD29-PE, clone TS2/16 (Biolegend, San Diego, CA, USA) to the desired concentration and incubated for 30 minutes at 4°C in the dark. The cells were washed with PBS+2% FBS and then re-suspended in 0.5 mL of PBS+2% FBS. All flow cytometry was performed on a BD LSR II flow cytometer (BD Biosciences, Europe).

### Perturbation experiments

Cells were incubated with an anti-CD11a monoclonal antibody clone TS1/22 (Thermo Fisher Scientific, Waltham, MA, USA) or a recombinant monoclonal antibody to integrin alpha 4 (CD49) clone Natalizumab (Absolute Antibody, Boston, MA, USA) for 10 min at 37°C. Cells were first seeded in channels with RPMI medium and were incubated for 10 min at 37°C to allow adhesion before starting the experiment.

## Acknowledgments

The project leading to this publication has received funding from the ANR grant RECRUTE, LABEX INFORM, the Région PACA, Institute CENTURI, Excellence Initiative of Aix-Marseille University – A*MIDEX, a French “Investissements d’Avenir” programme, Nanolane company and Alveole company. We thank the France Bioimaging Platform, funded by the French Agence Nationale de la Recherche (ANR–-10–-INBS–-04–-01, «Investments for the future»). We are grateful to Alphée Michelot for his support and advices with TIRF-FRAP imaging, to Laurence Borge for assistance with the use of the Cell Culture Platform facility (Luminy TPR2-INSERM).

## Supplementary files

### Supplementary movies

Movie 1: T cell motility phenotype under flow (8 dyn.cm^---2^) on ICAM-1 (left) and VCAM-1 (right) substrates. Bright-field images with magnification x10.

Movie 2: T cell motility phenotype under flow (8 dyn.cm^-2^) on mixed ICAM-1/ VCAM-1 substrates. Transmission images with magnification x10.

Movie 3: T cell motility phenotype under flow (4 dyn.cm^-2^) on mixed 50 % ICAM-1 / 50 % VCAM-1 substrate. Transmission (left) and RICM (right) images with magnification x63.

Movie 4: T cell motility phenotype under flow (8 dyn.cm^-2^) on ICAM-1 (left) and VCAM-1 (right) substrates. Top: Transmission images with magnification x63. Bottom: RICM images with magnification x63.

Movie 5: Flow triggers no calcium signaling. Relative intracellular calcium levels in crawling lymphocytes without and with flow, on ICAM-1 then VCAM-1 surfaces. Last part of the movie is the control experiment with ionomycin.

Movie 6: The level of integrins expression dictates orientation decision. First part of the movie (left) shows that on a mixed substrate 75% ICAM-1 – 25% VCAM-1, cells are mainly crawling upstream. By adding blocking antibody against LFA-1 (TS1/22) and decreasing the ratio LFA-1/ VLA-4, cells are mainly crawling downstream (right). Second part of the movie (left) shows that on a mixed substrate 25% ICAM-1 – 75% VCAM-1, cells are mainly crawling downstream. By adding blocking antibody against VLA-4 (Natalizumab) and decreasing the ratio LFA-1/ VLA-4, cells are mainly crawling upstream (right).

### Supplementary Information

Supplementary Information 1: Quantification of ICAM-1 VCAM-1 amounts on substrates

Supplementary Information 2: Quantification of LFA-1 and VLA-4 expression on effector T cells

## Supplementary Information

### Supplementary Information 1: Quantification of ICAM-1 VCAM-1 amounts on substrates

Anti-human CD106-PE and anti-human CD54-PE (eBioscience by Thermo Fischer Scientific, Waltham, MA, USA) antibody were used for the quantification of substrates coatings with mixed ICAM-1 and VCAM-1. First we set up a bulk calibration curve by measuring the fluorescence intensity of 41 μm thick channels filled with antibody solutions at concentrations of 1.5, 3, 5 and 7 μg.mL^-1^. Channels were pre-treated with 1% Pluronic F127 (Sigma-Aldrich, St Louis, MO) fluorescence to limit adsorption of antibodies on surfaces. Channels were nevertheless rinsed with PBS and the residual fluorescent intensity corresponding to adsorbed antibodies on the surface was measured and then subtracted to the previous measurements. Fig S1-a shows that the final values are proportional to the volume concentration of antibody. The molar weight of an antibody being of 150 kDA, then 1 μg.mL of antibody corresponds to 4 molecules.μm^-3^ and assuming that the signal is given by the total number of molecules in the thin channel then the volume concentration can be turned in a surface concentration for a channel of height 41 μm (Fig S1-b). For each condition of substrate preparation with a given solution of mixed ICAM-1/VCAM-1, two samples were prepared and stained either with CD106-PE or with CD54-PE at 10 μg.mL^-1^ overnight at 4°C. Fluorescent images were taken the next day. The fluorescent intensity on either ICAM-1 or VCAM-1 channel minus the fluorescent intensity measured on the Protein A channel was then converted into surface density by comparison with the calibration curves. Figure S1-c shows that, linear variations of Fc-ICAM-1 and Fc-VCAM-1 adsorbed on substrates were obtained versus respective ratio of Fc-ICAM-1 and Fc-VCAM-1 in solution.

**Figure S 1:**
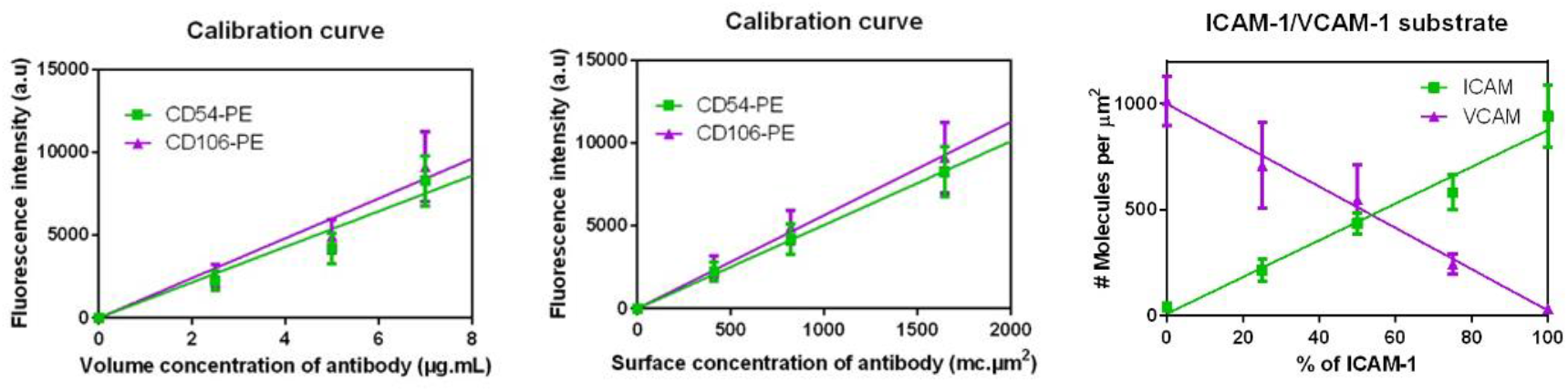
Quantification of substrates with mixed ICAM-1/VCAM-1. (A) Bulk calibration data obtained by measuring the fluorescence intensity of 41 μm thick channels filled with antibody solutions. (B) Conversion of bulk calibration data into a calibration curve linking fluorescence intensity with surface concentration of antibody. (C) Quantification of substrates coated with mixed ICAM-1/VCAM-1 versus the percentage of ICAM-1 in mixed ICAM-1/VCAM-1 solutions used for incubations. All data are mean + s.d, n= 3 independent experiments

### Supplementary Information 2: Quantification of LFA-1 and VLA-4 expression on effector T cells

Quantification LFA-1 and VLA-4 number per cell was performed by quantitative cytometry (Figure S 2) and yielded an average number per cell of 25000 for LFA-1 and 13000 for VLA-4.

**Figure S 2:**
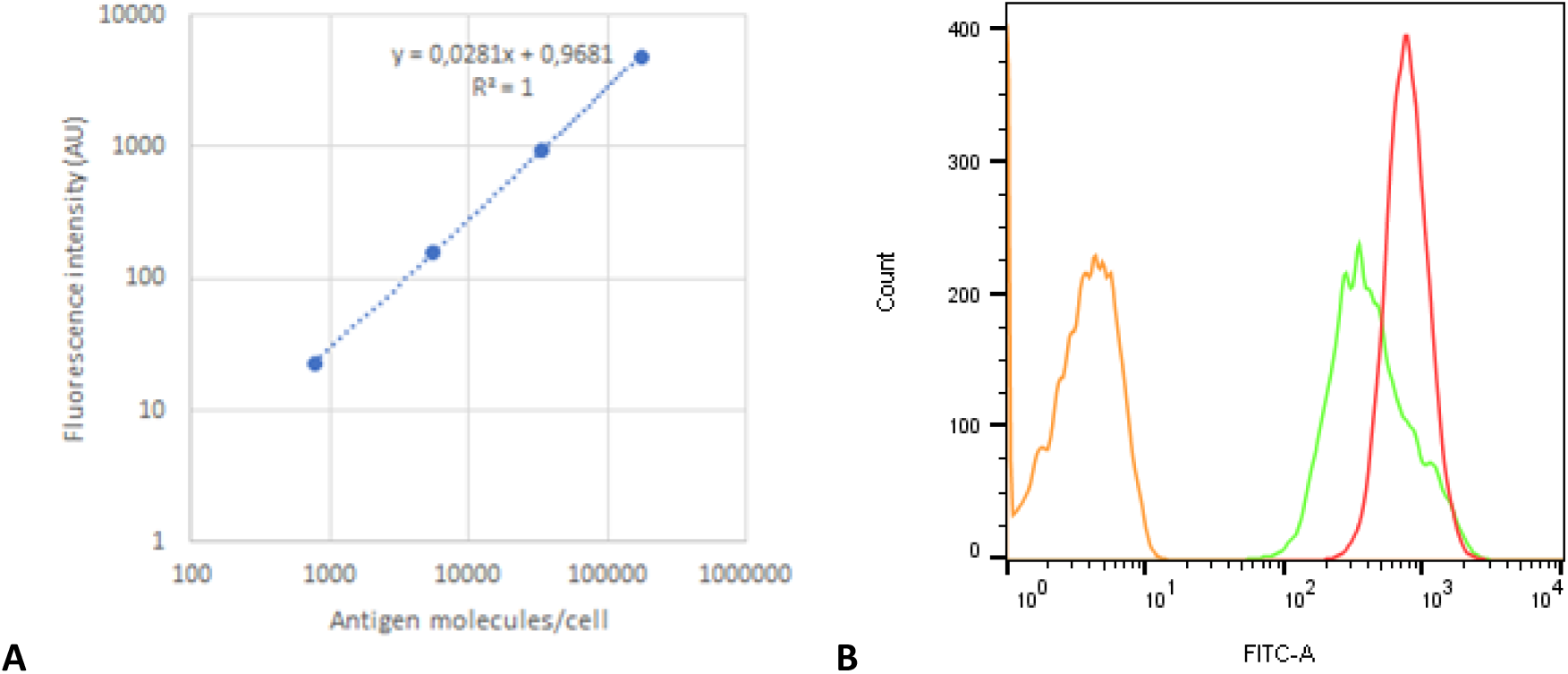
(A) Calibration curves with the secondary antibody and calibration beads (CellQuant calibrator kit, ref 7208, Biocytex) (B) fluorescence histograms of T cells stained by indirect immunofluorescence with specific monoclonal antibodies (CD49d (HP2/1) for VLA-4, green curve and CD11a (Hi111) for LFA-1, red curve).

